# The Symptomatology and Diagnosis of Domoic Acid Toxicosis in Stranded California Sea Lions: A Review and Evaluation of 20 Years of Cases to Guide Prognosis

**DOI:** 10.1101/2023.07.02.547442

**Authors:** Abby M. McClain, Cara L. Field, Tenaya A. Norris, Benny Borremans, Pádraig J. Duignan, Shawn P. Johnson, Sophie T. Whoriskey, Lorraine Thompson-Barbosa, Frances M. D. Gulland

## Abstract

Domoic acid (DA) is a glutaminergic excitatory neurotoxin that causes morbidity and mortality of California sea lions (*Zalophus californianus*; CSL) and other marine mammals due to a suite of effects mostly on the nervous and cardiac systems. Between 1998 and 2019, 11,737 live-stranded CSL were admitted to The Marine Mammal Center (TMMC; Sausalito, CA, USA), over 2,000 of which were intoxicated by DA. A plethora of clinical research has been performed over the past 20 years to characterize the range of toxic effects of DA exposure on CSLs, generating the largest dataset on the effects of natural exposure to this toxin in wildlife. Here we review published methods for diagnosing DA intoxication, clinical presentation and treatment of DA-intoxicated CSL, and present a practical, reproducible scoring system called the neuroscore (NS) to help assess whether a DA-affected CSL is fit for release to the wild following rehabilitation. The largest proportion of DA-intoxicated CSL were adult females (58.6%). The proportions of acute and chronic cases were 63.5% and 36.5%, respectively, with 44% of affected CSL released and 56% either dying naturally or euthanized. The average time in rehabilitation was 15.9 days (range 0-169) for all outcomes. Logistic regression models were used to assess the relationships between outcome (released vs. euthanized or died) and multiple variables to predict the outcome for a subset of 92 stranded CSLs. The best performing model (85% prediction accuracy; area under the curve = 0.90) consisted of four variables: final NS, change in NS over time, whether the animal began eating in rehabilitation, and the state of nutrition on admission. These results suggest that a behavioral scoring system is a useful tool to assess the fitness for release of stranded DA-intoxicated CSL.

## Introduction

Domoic acid (DA) is a potent glutaminergic excitatory neurotoxin that is produced by some species of marine diatoms in the genus *Pseudonitzschia*.^4,38^ The toxin accumulates in filter-feeding finfish, shellfish and other prey items which can cause toxicosis when consumed by vertebrates.^1,4^ Domoic acid toxicosis in humans was first recognized in 1987 when over 100 people became ill after consuming DA-contaminated mussels.^4,38^ Since then, many of the diverse effects of DA on mammals have been elucidated through experimental exposures of laboratory animals including zebrafish, rodents and non-human primates, and observations of naturally exposed marine wildlife.^15,18,20,26,33,37,44,45,48,54,55^ Amongst wildlife, most available information is derived from clinical and pathological studies of California sea lions (*Zalophus californianus*; CSL) that stranded along the California coast following natural exposure. Domoic acid intoxication was first documented in marine mammals in 1998 when over 400 CSL stranded along the California coast showing neurological signs and acute mortality.^18,43^ Since 1998, CSL have been intoxicated with DA annually, and a suite of characteristic clinical effects and histological lesions are consistently recognized. These lesions have also been described in southern sea otters (*Enhydra lutris nereis*) and northern fur seals (*Callorhinus ursinus*) stranded along the California coast.^27,31^

A decade after the first cases were recognized, a second clinical syndrome was described in CSL: chronic DA toxicosis. As opposed to acute toxicosis which immediately follows ingestion and absorption, chronic toxicosis is due either to repeated, low-level exposure of the toxin or to long-lasting effects of past sub-lethal exposure, as no toxin is present in the body at the time of clinical presentation.^6,15^ Acutely affected CSL typically strand in good body condition with neurological signs consisting of any combination of seizures, ataxia, head weaving, muscle tremors, decreased response to stimuli, and/or coma that either resolve within 7 days once DA is cleared from the body, or progress to a chronic epileptic state or mortality.^18^ The chronic syndrome is characterized by intermittent seizures and epilepsy associated with hippocampal atrophy, often accompanied by behavioral changes and disruptions of spatial memory.^15,48^ Neuronal necrosis, particularly in the ventral hippocampal complex, amygdala, pyriform lobe, and olfactory cortex is observed in acute cases, which progresses to hippocampal atrophy in the chronic syndrome.^6,45^ Degenerative cardiomyopathy attributed to direct action of DA on cardiac glutamate receptors has been described in both syndromes.^54^

Over recent years, the differentiation of these two syndromes has become increasingly difficult, presumably due to a combination of increased frequency of DA-producing harmful algal blooms in the California Current Ecosystem, and continued and/or repeated exposure of individual CSL.^29^ Following ingestion, DA can be detected in urine, serum, stomach contents, feces, amniotic fluid and milk. However DA is water soluble, has a half-life of only hours in blood and 24 hours in urine, and as the period from exposure to sample collection is often unknown, a negative result from tested body fluids does not rule out DA intoxication.^5,53^ Importantly, the prognosis differs between the acute and chronic syndromes; in acute cases symptoms may resolve, whereas in chronic cases they will not, resulting in poor prognosis for survival.^48^ Therefore, it is critical to define clinical signs and biomarkers that can be used as prognostic indicators in intoxicated animals outside of biological sampling, both for the management of CSL in rehabilitation and for best assessment of their fitness for release.^48,49^ Rapid determination of release potential allows for more humane management of animals, such as earlier euthanasia to relieve suffering of animals with little to no chance of survival post-release.

In this paper, we review the published methods for diagnosis of DA toxicosis in CSL as well as the clinical presentation and treatment of DA toxicosis in CSL based on published literature and 20 years of data collected by The Marine Mammal Center (TMMC), a rescue and rehabilitation facility for stranded marine mammals along central and northern California, based in Sausalito (CA, USA). To aid future clinical management of DA toxicosis in CSL, we present a behavioral assessment tool to aid clinicians in determining whether a CSL affected by DA is fit for release.

## Materials and Methods

### Literature Review

Search criteria included peer-reviewed journal articles, conference proceedings, internal case definitions, and book chapters that investigated methods for the diagnosis of DA toxicosis in CSL and laboratory animals. Additionally, the effects of DA toxicosis in CSL, laboratory animals, and other marine species as well as the clinical presentations of DA intoxication were also explored. A set time frame for publication date was not included in the search criteria, and relevant published literature was included regardless of publication date. Interrogated databases included Google Scholar and Pubmed as well as an internal list of DA-related publications and book chapters from TMMC. Key words used to search online databases included “domoic acid”, “sea lion”, “zalophus”, “sea otter”, “cardiac”, “in utero”, “neurologic”, “fur seal”, “marine mammal” and “ophthalmic”.

### Case definition

Inclusion criteria for antemortem diagnosis of DA toxicosis (acute or chronic) in CSLs were seizures, ataxia, head weaving, tremors, blindness, blepharospasm, reduced responsiveness, coma, and/or abnormal behavior that included disorientation, unusually human-oriented or inquisitive, or aggressive behavior, or repetitive behaviors (i.e., frantic pacing, swimming in tight circles, rocking on the pen floor, and/or chewing on body parts or objects) (TMMC case definition). A relative eosinophilia (greater than 0.68 × 10^9^/L), elevated hematocrit (median = 49.5%), and/or reduced serum cortisol concentration (<20mg/dL) on admission bloodwork provided further indication of DA intoxication (TMMC case definition).^13^

Antemortem differentiation between acute and chronic DA intoxication was based on stranding history, clinical signs, and imaging when available. Acute DA intoxication was presumed to occur when a group of ≥5 sea lions with neurological signs stranded within 48 hours and 80 km of each other and if available, documentation of an active *Pseudonitzschia* bloom or elevated DA levels from monitoring programs like the Southern California Coastal Ocean Observing System (SCCOOS) and the Central and Northern California Ocean Observing System (CeNCOOS). When available, post-mortem lesions in at least one of the sea lions consistent with acute DA toxicosis in the hippocampus provided further evidence for an acute DA intoxication event.^15^ Acute DA intoxication was usually inapparent on gross necropsy examination but was confirmed on histology by the presence of neuronal necrosis/sclerosis in the Cornu Ammonis of the hippocampus that may also affect neurons of the dentate gyrus, para-hippocampal gyrus, amygdaloid body, olfactory and pyriform lobe cortex.^39,45^

Inclusion criteria for chronic DA intoxication were defined based on stranding history in which a single animal stranded in an abnormal location (delta, urban area, etc.), typically in poor body condition, and as an isolated event with neurologic abnormalities.^15^ Antemortem magnetic resonance imaging (MRI) showing hippocampal atrophy provided further evidence of chronic DA intoxication when available. Postmortem histological diagnosis of chronic DA intoxication was based on often asymmetric atrophy of the hippocampal complex characterized histologically by neuronal loss and gliosis.^39,45^

### Treatment

Therapy was aimed at control of symptoms as previously described in the literature and the TMMC pharmacopeia.^18^ Phenobarbital (Golden Gate Pharmacy, Novato, CA 94949, USA) was administered for seizure control during rehabilitation and was started upon the initial diagnosis of DA toxicosis (4 mg/kg IM q12h for two days followed by 2mg/kg q12h for five more days (IM or PO if eating). Lorazepam (Golden Gate Pharmacy, Novato, CA 94949, USA), midazolam (Almaject, Morristown, NJ 07690, USA), and/or diazepam (Hospira, Lake Forest, Il 60045, USA) were used in conjunction with phenobarbital for the treatment of active, break-through seizures (0.2mg/kg IM PRN).

Adjunct therapy to help minimize inflammation and cellular necrosis included an antioxidant, alpha lipoic acid (10mg/kg SQ q24h; Golden Gate Pharmacy, Novato, CA 94949, USA) ^12^ and an anti-inflammatory dose of dexamethasone (Bimeda-MTC Animal Health Inc, Cambridge, Ontario N3C 2W4, Canada) or prednisone (Novitium Pharma LLC, East Windsor, NJ 08520, USA) unless contraindicated due to co-morbidities such as corneal ulceration. Supportive subcutaneous fluids (lactated Ringer’s solution, LRS, Vetivex, Dechra Veterinary Products, Overland Park, Kansas 66211 USA; normosal-R, Plasma-Lyte A, Abbott Animal Health, Abbott Park, Illinois 60064, USA) were administered to anorexic animals at 20-25ml/kg/day for 3-5 days or until animals started eating.

As DA is retained in fetal amniotic and allantoic fluids leading to prolonged persistence in circulation with subsequent progressive disease and fetal developmental abnormalities,^16,26^ abortion was induced in pregnant females by administering dexamethasone at 0.25mg/kg IM q24h for three days. If abortion did not occur by day three, a single 500 microgram dose of prostaglandin F2alpha (Lutalyse®, Zoetis, Parsippany-Troy Hills, NJ, 07054, USA) was administered. Prophylactic ceftiofur crystalline free acid (Excede, Zoetis, Parsippany, New Jersey 07054, USA) was administered IM (6.6mg/kg) to reduce risk of pyometra in debilitated animals.

### Study Design

A retrospective analysis of the clinical presentation, demographics, and survival to release of 2,447 CSL diagnosed with DA intoxication and admitted to TMMC between 1998-2019 was performed. A subset of 92 CSL affected by DA and admitted to TMMC between July 2015 and August 2017 was analyzed to determine the performance of a quantitative behavioral assessment tool, referred to as the neuroscore (NS), in predicting whether a CSL affected by DA was a candidate for release. The subset of CSL were chosen because they all had at least 3 complete NS performed with the most up-to-date NS and criteria used consistently by experienced TMMC veterinarians. The NS criteria has been adapted over time based on the changing symptomatology of DA intoxication. Stranding data and clinical presentations, a review of the current diagnostic tests available to differentiate and diagnose acute and chronic DA intoxication, and the performance of the NS in helping to predict whether a CSL is fit for release as well as its limitations are reported.

### Clinical assessments

All animals stranded along the California coast between approximately 34°22’23” N and 40°0’4” N and were transported to TMMC for assessment and possible rehabilitation. Physical examinations were performed on all animals upon admission under manual or chemical restraint. Age class was estimated based on body mass, straight body length, tooth development, and development of the sagittal crest in males.^17^ State of nutrition was estimated based on a body condition scale of 1 to 5, with 1 being severely malnourished and 5 being over-conditioned (TMMC internal scale). Blood samples were collected via the caudal gluteal vein directly into vacutainer collection tubes and complete blood count and serum chemistry were analyzed as previously published.^18,50^ Urine was collected when possible via cystocentesis or catheterization, and abdominal ultrasound examination performed on female CSL to assess for pregnancy. MRI was performed under general anesthesia using published anesthetic protocols.^8^ Sea lions were housed with one or more conspecifics of similar age, in either dry pens (if non-responsive initially), or pens with access to a saltwater pool and concrete deck space. They were offered frozen, thawed herring two to three times daily and records indicating whether the animal was eating or not were maintained. Sea lions were diagnosed with domoic acid intoxication using the earlier described inclusion criteria.

### Release Assessment

A quantitative behavioral assessment tool (NS) aimed at quantifying the severity of DA-associated effects on stranded CSL was developed at TMMC and was performed on CSL diagnosed with DA intoxication as a standardized method to assess the clinical response to treatment. This assessment is similar to the method previously reported,^18^ although the assessment has been adjusted over the years based on the changing symptomatology (acute vs. chronic) observed in CSL. The NS (Figure 1) was modeled after pain scoring systems in human neonates such as the premature infant pain profile and the neonatal infant pain scale,^24,47^ and is a sum of multiple components assessing posture, mentation, the presence of abnormal movements, response to auditory and tactile stimuli, and locomotor skills. The sum of these components is the NS, where a lower NS is consistent with an animal with a higher probability of survival after release (as determined by low likelihood of re-stranding). The cutoff value used at TMMC to determine whether a CSL was a candidate for release was a NS of 10 or below. The NS assessments began two days after an animal finished the previously described course of treatment (i.e., two days after the final dose of phenobarbital) to minimize potential residual sedative effects following phenobarbital administration. A single NS assessment was performed every other day by experienced TMMC clinical veterinarians until three scores were obtained, upon which the NS trend as well as the overall clinical progression of the animal were taken into consideration to determine whether to release the animal.

**Figure 1:**
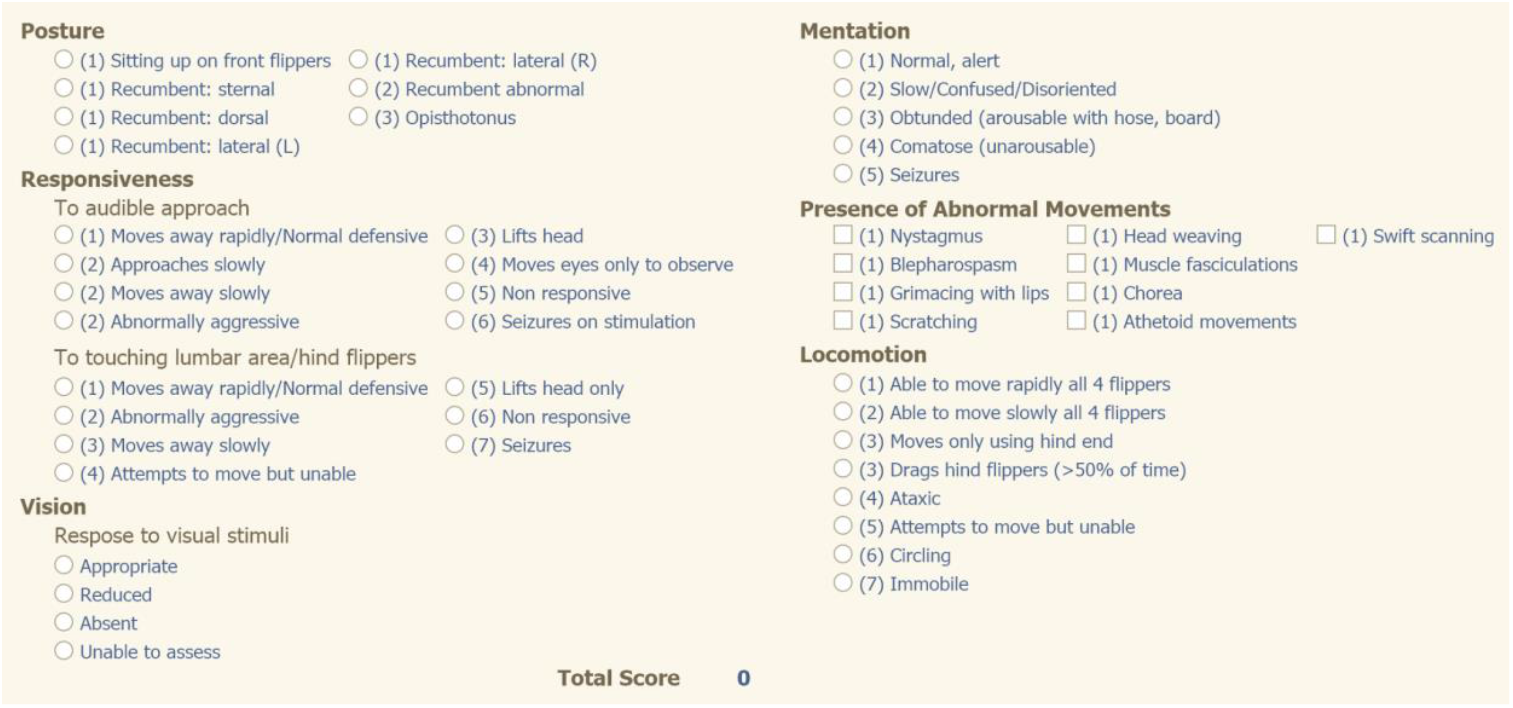
Neuroscore criteria used at The Marine Mammal Center to assess the clinical responsiveness (or the clinical response to treatment) of stranded California sea lions intoxicated by domoic acid. Circles next to terms indicate that one option should be selected, squares indicate more than one option may be selected. The assessor chooses a single score based on the animal under assessment for the sections “Posture”, “Mentation”, “Responsiveness”, and “Locomotion” and marks all that apply for the “Presence of Abnormal Movements” section. An animal that receives a score of 10 or less is considered a candidate for release.

### Statistical Analysis

Differences in the proportion of CSL stranding due to DA between age classes, sex, outcome (released vs. died/euthanized) and in the proportion of acute vs. chronic cases were tested using Pearson’s chi-squared test on the full dataset (CSL stranding between 1998-2019). Correlations between variables were tested using generalized linear models and likelihood ratio tests, with model specifics provided in the results section.

We tested whether it was possible to use different variables to predict the outcome (release vs. death) after stranding due to DA on the subset of 92 CSL stranding between 2015-2017. Candidate variables included the state of nutrition, straight body length, presence of a circulating eosinophilia, and diagnosed co-morbidities at admission, and whether the animal began eating while in rehabilitation, in conjunction with the NS results. We report which combination of variables were the best predictors of releasability to aid rehabilitation facilities in determining whether a DA-affected CSL is fit for release (as determined by low likelihood of re-stranding). Generalized linear models with logit link function and binomial error distribution were constructed for different combinations of candidate variables, and their performance was compared using 10-fold cross-validation error, AIC and AUC (area under the curve).^11^ Cross-validation was used to explicitly test the predictive performance of a model.^19^ For the 10-fold cross-validation test, the dataset is randomly divided into 10 parts. For each of those 10 parts in turn, a model is fit using the 9 parts (training data) not included in the current part, after which the fitted model is used to predict the outcome of the data in the current part (test data). This approach repeatedly simulates a situation where a model is constructed using observed data and is then used to predict the outcome of future observations that have not been used to construct the model. Statistical results were reported in terms of level of statistical support for a tested correlation or difference between models, as opposed to a binary “significant or not significant” classification based on an arbitrary P value cutoff of 0.05. “No support” is used when P values are high (around 0.1 and above), “weak support” when P values are close to 0.05 (around 0.01-0.1), “moderate support” when P values are small (around 0.001-0.01), and “strong support” when P values are very small (roughly < 0.001). This reporting method emphasizes the fact that statistical significance is determined on a continuum, and follows current reporting recommendations.^34^

Analyses, data manipulation and plotting were performed in R, using packages lme4, ggplot2, pROC and dplyr.^3,40,41,51,52^

## Results

### Review of all Diagnostic Testing

A wide variety of tests have been evaluated to investigate and characterize the effects of DA toxicosis in CSL as alternatives to detection of DA itself in body fluids or tissues, but few have been evaluated for sensitivity or specificity or have been applied to the clinical setting on a regular basis. Changes in circulating numbers of eosinophils, serum apolipoprotein E and cerebrospinal fluid proteins have been detected in groups of sea lions with DA toxicosis compared to unaffected animals; and DA antibodies have been detected in sea lions with chronic toxicosis.^13,25,35,36^ Antemortem MRI evaluation of hippocampal size^6,32^ and non-invasive diagnostic methods have also been explored. Behavioral studies in CSL affected by DA found a direct correlation between long-term spatial memory and right-sided hippocampal structure.^9^ Measurement of habituation rates to an auditory stimulus and the presence of persistent abnormal behaviors such as head weaving, muscle fasciculations, etc., can also help diagnose DA toxicosis.^7,53^ As neurons in the olfactory bulb may also undergo DA-induced necrosis, olfactory testing has also been evaluated for the diagnosis of DA intoxication.^28^ A summary of these tests, their results, and potential limitations are listed by publication date in Table 1.

**Table 1:** Summary table of published methods to aid in the diagnosis of acute domoic acid toxicosis, chronic domoic acid toxicosis or both. DA=domoic acid; CSL=California sea lion; EEG=electroencephalogram; MALDI-TOF=matrix-assisted laser desorption/ionization-time of flight; MRI=magnetic resonance imaging.

| <u>Test/Study</u> | <u>Number of CSL in study</u> | <u>Findings</u> | <u>Limitations</u> | <u>References</u> |
| --- | --- | --- | --- | --- |
| EEG | 29 | Numerous epileptiform discharges were observed in chronically affected CSL; affected areas were primarily multifocal; discharges consisted of short duration, high-voltage spikes or sharp waves and intermittent rhythmic delta activity often followed by brief background attenuation; features were localized to the posterior aspects of both hemispheres with voltage maxima in one hemisphere or the other | Requires heavy sedation or general anesthesia; need access to an EEG machine as well as knowledge about how to interpret results; may be less effective in diagnosis acute DA intoxication | Goldstein et al. 2008 |
| Circulating eosinophilia detected by complete blood count | 39 | The median eosinophil count in the DA group was $0.925 \times 10^9$ eosinophils/L, which was significantly higher than the non-DA group; presence of an eosinophilia may be a cost-effective biomarker for DA exposure | Blood results were combined instead of tested individually, so not all DA affected CSL may have an eosinophilia; does not differentiate between acute and chronic exposures; need a blood sample | Gulland et al. 2009 |
| Auditory Habituation rate | 39 | CSL with DA toxicosis took significantly more exposures to an auditory stimulus to habituate in the first test phase than did controls; a pre-set habituation measure correctly identified 50% of CSL with DA toxicosis and falsely diagnosed 7% of control CSL | Need a quiet space with minimal distractions to conduct the test | Cook et al. 2011 |
| Anti-DA antibodies | 22 (17 cases; 5 controls) | 11 of 17 CSL cases had detectable levels of anti-DA-specific antibodies in their serum by indirect ELISA indicating previous exposure to DA induces a detectable antibody response; in the future, could be a point-of-care test | Not clinically available; diagnoses previous exposure but not extent of exposure including duration and persistence; need a blood sample | Lefebvre et al. 2012 |
| MRI | 53 | Hippocampal volumes were less in CSL with chronic DA intoxication than non-DA intoxicated CSL; chronically affected CSL showed thinning of the parahippocampal gyri compared to acute DA and non-DA CSL; all CSL with chronic DA exhibited structural damage to the limbic system and always had hippocampal atrophy | Requires prolonged general anesthesia and access to an MRI machine; increased cost and resources | Montie et al. 2012 |
| Plasma Proteomics | 32 (Neely 2015)<br>107 (Neely 2012) | No single peptide peak from MALDI-TOF profiling was a good indicator of acute DA toxicosis; neural networks trained using 104 peptide peaks had a sensitivity of 100% and specificity of 60% for diagnosis of acute DA toxicosis. Different charge forms of apolipoprotein E along with the eosinophil count can help diagnose chronic DA toxicosis. | Neither test has been translated into the clinical setting; highly technical and need a specialized laboratory to run and analyze results; need a blood sample | Neely et al., 2012. Neely et al. 2015 |
| Cerebral spinal fluid proteomics (CSF) | 11 | CSF protein concentration was elevated in CSL with DA toxicosis suggesting a more permeable or less selective blood brain barrier; proteins were not identified as biomarkers, but concentration increased | Requires general anesthesia and increased technical skill to obtain samples; not translated into the clinical setting; need a specialized laboratory for equipment and to analyze results | Neely et al. 2015 |
| Behavioral changes | 169 | Described 23 abnormal behaviors classified according to posture, movement, and ingestion abnormalities; head weaving and muscle fasciculations were observed for significantly longer durations in DA animals compared to controls; swift scanning was observed in the DA group but not the non-DA group; dragging hind flippers was observed in the DA group but not the non-DA group; presence and duration of abnormal behaviors resulted in a diagnostic accuracy rate of 88% and false positive likelihood of 15% | Need a quiet space with minimal distractions for monitoring; ideally need a blind or other structure to conceal the observer | Wittmaack et al. 2015 |
| Olfaction | 55 | Control CSL spent significantly more time with a scented object than an unscented object; chronically affected DA CSL showed significantly higher variance in proportion of time spent with the scented object vs the unscented object | Need space to set up and perform the test; test does not assess olfactory abilities across a range of ecologically valid contexts (i.e., mother-offspring recognition) | De Maio et al. 2018 |

Despite the variety of antemortem tests that have shown promising results as diagnostic tools to help characterize effects of DA intoxication, not all have been applied clinically on a larger scale either due to logistical reasons, test and reagent development and/or availability, lack of resources, or low specificity. In addition to diagnostic tools, inexpensive, practical, and minimally invasive methods to aid clinicians in determining whether a CSL affected by DA is clinically fit for release are still needed.

### Clinical Presentation and Diagnostics

Various clinical presentations previously described in the literature were observed in CSL with DA intoxication including, but not limited to, head weaving, ataxia, seizures, and coma that varied in severity depending on the animal. Cardiomyopathy, blindness and the presence of an eosinophilia (diagnosed as an absolute eosinophil count greater than 0.68 × 10^9^/L) have also been well-documented in the literature.^13,15,45,54^ Cardiomyopathy and blindness are thought to be caused by DA binding to glutamate receptors on cardiac muscle fibers and to retinal neurons, whereas the cause of an eosinophilia is unknown but is hypothesized to be related either to adrenal gland function or caused by exposure to DA itself independent of adrenal gland function.^13,14,45,54^ Historically, an eosinophilia was reported in both acute and chronically intoxicated CSL in the year 2009 with a mean of 0.925 × 10^9^/L.^13^ Similarly, we found this trend to be consistent in our full dataset of stranded CSL between 1998-2019. Data differentiating between acutely intoxicated CSL and chronically intoxicated CSL were available on 1640 out of the 2448 CSL within the full dataset. Of these 1640 CSL, eosinophil data was available on 642, where 63.91% (n = 540) had an eosinophilia on admit blood work; 70.6% (n=293) of acutely intoxicated CSL and 61.7% (n=140) of chronically intoxicated CSL had an eosinophilia. We found a slightly higher mean eosinophil count of 1.07 × 10^9^/L (acute and chronically intoxicated CSL combined) compared to previously published data.

As DA-producing blooms are most common in spring and early summer, it is not uncommon for late pregnancy CSL to strand with DA intoxication.^26^ To date, clinical treatment has been aimed at saving the adult rather than the fetus, as fetal damage due to in utero exposure is suspected, making offspring prognosis poor.^26,44^ The number and outcomes of all CSL diagnosed as pregnant, blind, having an eosinophilia on admission blood work or with cardiomyopathy between 1998 and 2019 are reported in Table 3.

### Overall Trends

Between 1998 and 2019, TMMC rescued a total of 11,737 CSL; 21% (n = 2,447) of which were diagnosed with DA toxicosis (Figure 2). Strong statistical support for an increase in CSL stranding due to DA intoxication over time was found (generalized linear model with log link function and Poisson error distribution; ChiSq = 41.7, p-value <0.001) where, on average, the number of DA-associated strandings increased by 2.1% each year, which may in part be due to an increase in the total CSL population over time.^23^ Of the DA-associated strandings, the proportions diagnosed with acute and chronic syndromes were 63.5% (n = 1042) and 36.5% (n = 598), respectively, with strong statistical support for an annual increase of 7.4% of the average proportion of chronic DA strandings (ChiSq = 42.3, df = 1, p-value < 0.0001). We were unable to categorize a total of 807 cases due to the lack of use of the classification of acute or chronic DA intoxication at admission by the clinician.

**Figure 2:**
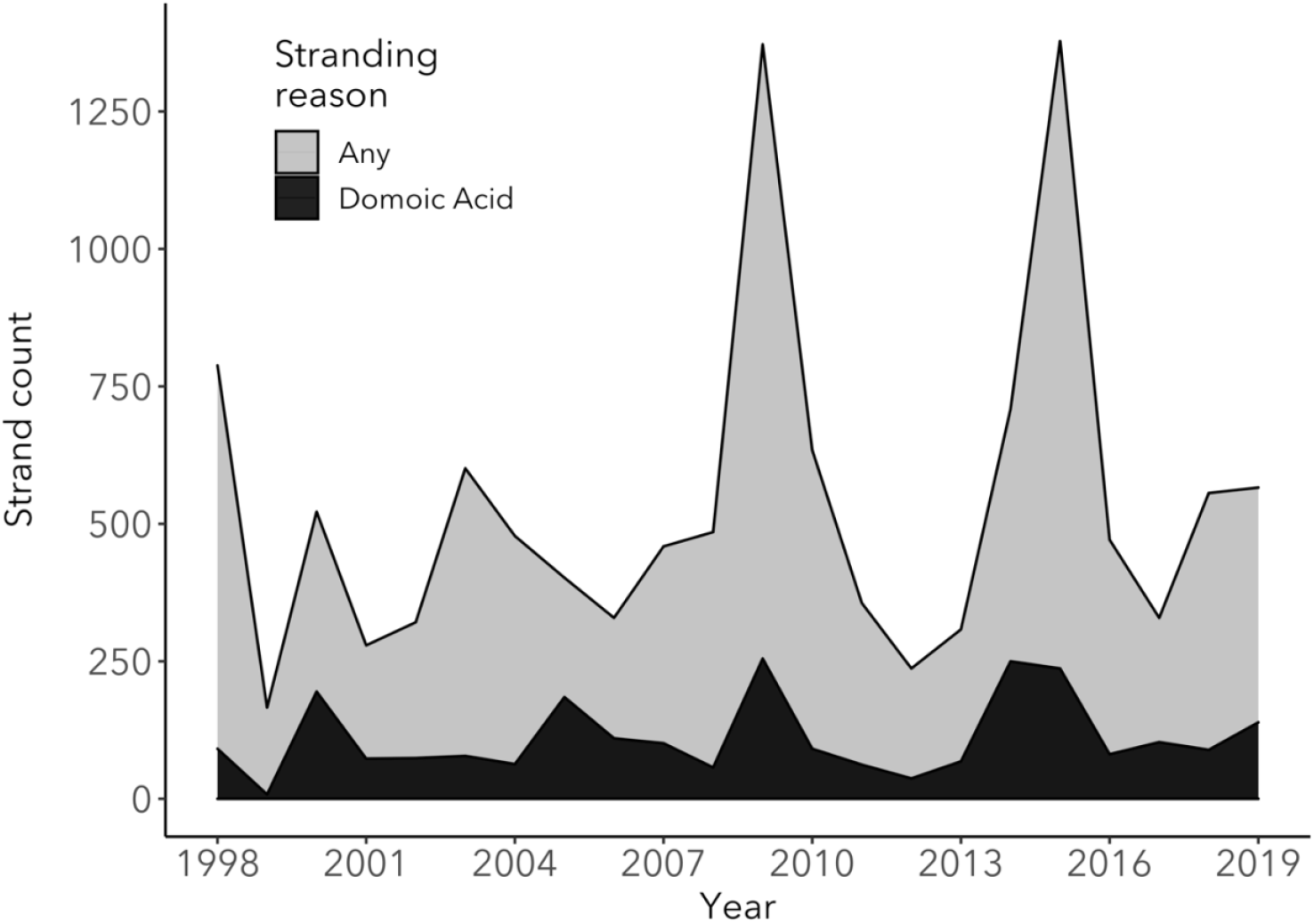
Total number of stranded California sea lions at The Marine Mammal Center (TMMC) due to any cause (light gray) compared to the number of California sea lions stranding at TMMC due to domoic acid intoxication (black) between 1998 and 2019.

Between 1998 and 2019, 44% (n = 1,074) of stranded CSL with DA intoxication (acute or chronic) were released, while 56% (n = 1,366) died or were euthanized. Of the animals that died or were euthanized, 42.4% (n = 579) died naturally and 57.6% (n = 787) were euthanized. A total of 149 animals with DA intoxication re-stranded (6.1%) after being released, a rate that was higher than the re-strand rate due to causes of stranding excluding DA intoxication during this same period (3.7%). This rate considers each re-strand as a separate event even if the same animal re-stranded multiple times.

Strong statistical support was found for age and sex effects (ChiSq = 3698, df = 4, p-value <0.001; ChiSq = 881, df = 1, p-value <0.001, respectively). Most sea lions with DA toxicosis were adult female, followed by juvenile males, subadult males, subadult females, yearling females, and adult males (Table 2). The average rehabilitation times (number of days between admit and disposition date) were 24.9 days (range = 2-169) for animals that were released (excluding the animals that were placed in permanent human care), 8.3 days (range = 0-98) for those that died or were euthanized, with a combined (survived or died) average of 15.9 days for all outcomes.

**Table 2:** Summary of California sea lions diagnosed as pregnant, blind, or with cardiomyopathy as well as diagnostics performed and the percent of sea lions that survived. All California sea lions were intoxicated by domoic acid between 1998 and 2019 and were rescued by The Marine Mammal Center. DA=domoic acid, CSL=California sea lion.

| <u>Clinical Presentation</u> | <u>Number of CSL</u><br><u>(% of DA intoxicated</u><br><u>stranded CSL)</u> | <u>How Diagnosed</u> | <u>Findings</u> | <u>% Adults survived</u> |
| --- | --- | --- | --- | --- |
| Pregnant | 82 (5.7 of adult females) | Ultrasound | Viable or non-viable fetus | 56.1 |
| Blind | 27 (1.1) | Physical examination | Non-visual | 3.7 |
| Cardiomyopathy | 53 (2.2) | Auscultation, echocardiography, electrocardiography, and/or histopathology | Echocardiography: decreased fractional shortening, decreased ejection fraction, decreased cardiac output, valvular insufficiency, abnormal septal wall movement.<br>Electrocardiography: First and/or third- degree<br>atrioventricular<br>block, premature<br>atrial and ventricular<br>complexes. <sup>2</sup><br>Auscultation:<br>murmurs auscultated | 13.2 |
| Circulatory<br>eosinophilia | 356 (68.7) | Hematology at<br>admission | Absolute eosinophil<br>count greater than<br>$0.68 \times 10^9/L$ | 66.7 |

**Table 3:**
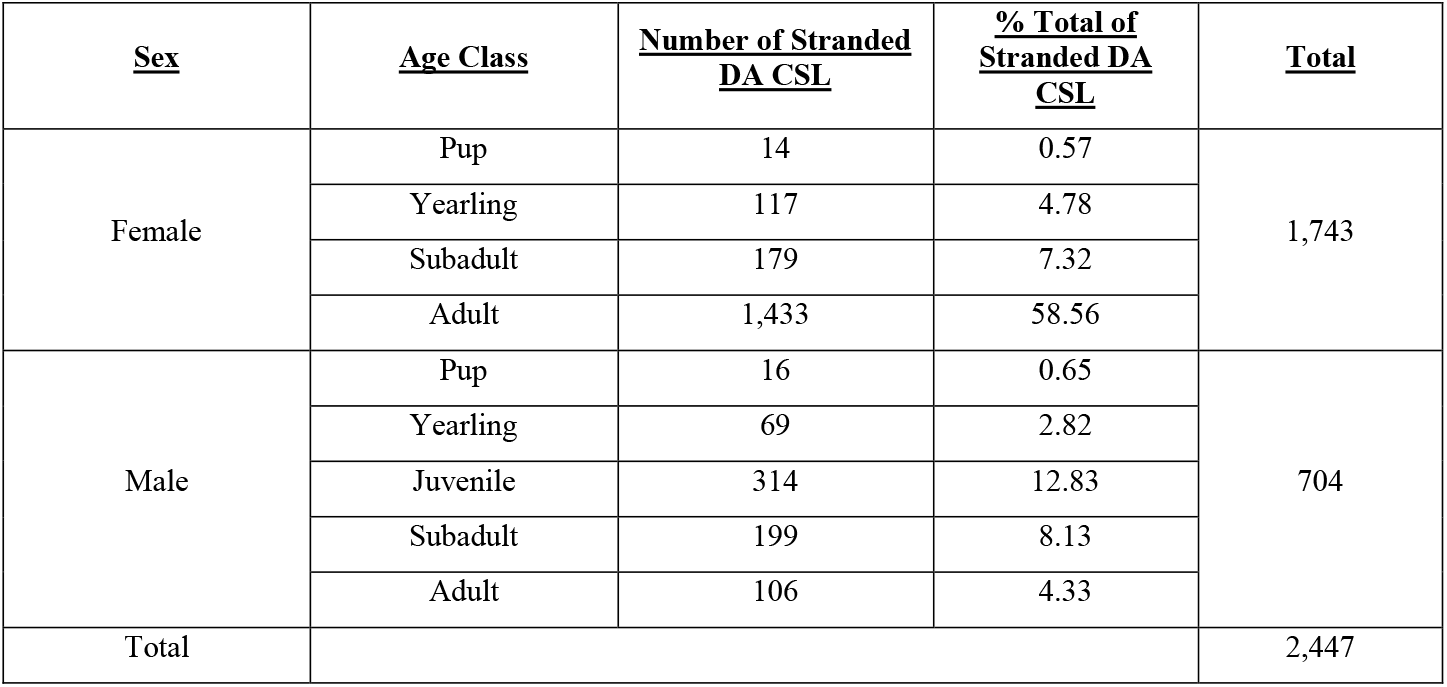
Summary of age class and sex distribution of stranded California sea lions diagnosed with domoic acid intoxication (acute, chronic, or acute on chronic) between 1998 and 2019 at The Marine Mammal Center.

| <u>Sex</u> | <u>Age Class</u> | <u>Number of Stranded DA CSL</u> | <u>% Total of Stranded DA CSL</u> | <u>Total</u> |
| --- | --- | --- | --- | --- |
| Female | Pup | 14 | 0.57 | 1,743 |
|  | Yearling | 117 | 4.78 |  |
|  | Subadult | 179 | 7.32 |  |
|  | Adult | 1,433 | 58.56 |  |
| Male | Pup | 16 | 0.65 | 704 |
|  | Yearling | 69 | 2.82 |  |
|  | Juvenile | 314 | 12.83 |  |
|  | Subadult | 199 | 8.13 |  |
|  | Adult | 106 | 4.33 |  |
| Total |  |  |  | 2,447 |

### Release Assessment

Records from a subset of 92 DA intoxicated CSL that stranded between July 2015 and August 2017 were reviewed to assess the predictive accuracy of the NS in determining whether a DA-affected CSL was fit for release. Other potential candidate variables that aid in determining releasability, and components of the NS which were most important in determining whether a DA-intoxicated CSL was fit for release, also were assessed. Table 4 lists all models tested along with their results. Of note, the best predictors of releasability were the third NS, whether the animal started eating while in rehabilitation, the difference in NS over time with a decreasing trend of NS values, and the state of nutrition at admission. Utilizing these four variables, the model correctly predicted whether a CSL was released or not 85% (AUC = 0.90) of the time. The presence of eosinophilia, co-morbidities, and the admission straight body length had no significant correlation with the outcome of the animal and thus were not included in the models. Of the 92 animals in the subset, 46.7% died or were euthanized (n=43) and 53.2% were released (n=49). Nine animals that died or were euthanized had full histopathology collected on necropsy and eight of the nine animals had brain and/or cardiac lesions consistent with domoic acid toxicity ranging from unilateral to bilateral hippocampal atrophy, inflammation, gliosis, and/or neuronal degeneration; myocardial degeneration, necrosis, or edema. The animal that did not have lesions consistent with domoic acid toxicosis was diagnosed with meningitis. ^49^ The remaining animals that died or were euthanized either did not have histopathology collected or did not have full histopathology collected to include the brain or heart.

**Table 4:**
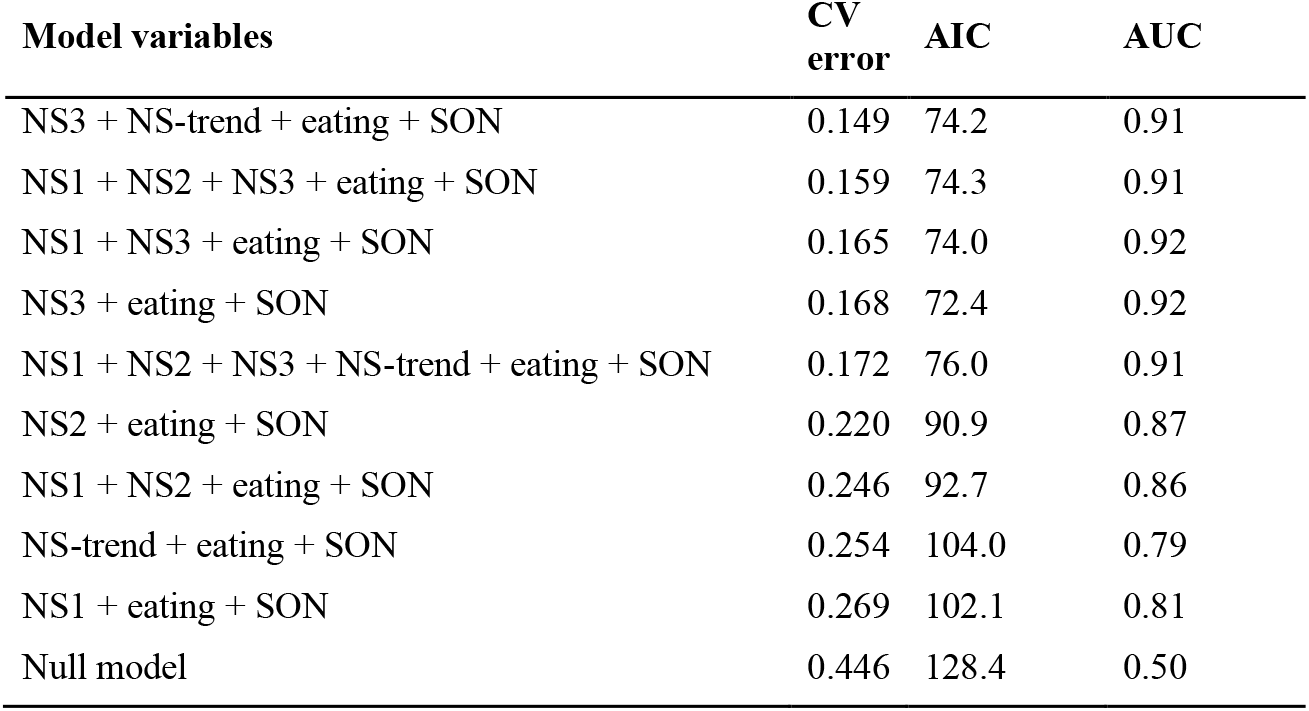
Performance of the top models predicting outcome (release vs. death) using different combinations of variables. CV error = 10-fold cross-validation error; AIC = Akaike Information Criterion; AUC = area under the curve. The models are ranked according to CV error, which should be most closely correlated with predictability. NS1, NS2 and NS3 = neuroscores at day 2, 4 and 6 after completion of phenobarbital; NS-trend = the slope of the regression between time and NS; eating = started eating while in rehabilitation; SON = state of nutrition at admission. Null model = model without variables, for comparison of the different statistics.

| Model variables | CV error | AIC | AUC |
| --- | --- | --- | --- |
| NS3 + NS-trend + eating + SON | 0.149 | 74.2 | 0.91 |
| NS1 + NS2 + NS3 + eating + SON | 0.159 | 74.3 | 0.91 |
| NS1 + NS3 + eating + SON | 0.165 | 74.0 | 0.92 |
| NS3 + eating + SON | 0.168 | 72.4 | 0.92 |
| NS1 + NS2 + NS3 + NS-trend + eating + SON | 0.172 | 76.0 | 0.91 |
| NS2 + eating + SON | 0.220 | 90.9 | 0.87 |
| NS1 + NS2 + eating + SON | 0.246 | 92.7 | 0.86 |
| NS-trend + eating + SON | 0.254 | 104.0 | 0.79 |
| NS1 + eating + SON | 0.269 | 102.1 | 0.81 |
| Null model | 0.446 | 128.4 | 0.50 |

We also assessed which components of the NS were the most important in determining fitness for release. In a situation where all combinations of the different NS subsections were possible (classification tree modeling), auditory approach response, tactile response, and mentation were the most important. When using only one subsection at a time in a logistic regression model, the subsections that correlated most strongly with outcome were auditory approach response, tactile response, and locomotion.

As the survivability of released CSL with DA intoxication is unknown and outside the scope of this study, the re-strand rate was used as an assessment of the effectiveness of the NS in aiding clinicians in determining whether an intoxicated CSL was fit for release. At TMMC, a NS of 10 for the third/final NS performed has been used since 2006 as a cut-off to differentiate between a releasable and non-releasable animal. Prior to the implementation of the NS in the release assessment (1998 – 2006), the average re-strand rate for DA intoxicated CSL was 8.2%. In the time frame after the consistent use of the NS during the release assessment (2007 – 2019), the average re-strand rate was 6%. The differences between the re-strand rates were not statistically supported, however the consistent implementation of the NS during the release assessment was associated with an improved average re-strand rate for released DA intoxicated CSL.

## Discussion/Future Directions

Despite the extensive research and effort applied to further our understanding of DA intoxication in marine wildlife, humans, and in laboratory animal models over the past 20 years, the clinical management of CSL and other marine mammals diagnosed with DA toxicosis remains challenging. Rehabilitation facilities have the responsibility to rehabilitate and release stranded marine mammals with a high “likelihood of survival” post-release as per NOAA guidelines.^49^ We reviewed published literature to summarize diagnostic tests used to help detect the spectrum of lesions and effects associated with the acute and chronic syndromes as well as treatment strategies for CSL intoxicated with DA, presented stranding statistics of DA-affected CSL over the past 20 years and evaluated a promising tool with candidate variables to aid rehabilitation facilities in determining whether a DA-affected CSL is fit for release (as determined by high likelihood of survival and low likelihood of re-stranding).

Of the previously published diagnostic tests listed in Table 1, the presence of an eosinophilia was the only previously published test evaluated in this study for correlation with “being released” with a low likelihood of re-stranding, and no correlation was found. Thus, the presence of an eosinophilia on admission was not included in the models. To date, clinical signs, especially when observed in CSL that strand in spatial and temporal clusters, are the most useful initial guide for a clinician that a CSL has DA toxicosis. An eosinophilia on blood work should increase the clinician’s suspicion of DA intoxication, especially as there are multiple other diseases that mimic DA toxicosis, such as bacterial meningitis or protozoal encephalopathy. CSF proteomics have promise for aiding clinicians in diagnosing DA exposure as multiple CSF proteins were found to be elevated in DA-intoxicated CSL compared to control CSL,^36^ and presence of DA antibodies shows promise as 11 of 17 CSL chronic DA cases had antibodies.^24^ A definitive diagnosis of hippocampal atrophy can be obtained with MRI. Using the entire clinical picture of the animal, the state of nutrition at admission, the third and final NS, the change in NS over time, and whether the animal started eating while in rehabilitation can help guide clinicians on the decision to release an animal with DA intoxication if MRI is not possible.

To date, research has focused on the diagnosis and pathological effects of DA toxicosis in mature CSL. Over the past 20 years, 30 stranded pups were also diagnosed with DA intoxication at TMMC, and some affected pregnant females that do not strand may give birth to live neonates. The ability of DA to cross the placenta and recirculate between the fetus and amniotic fluid, as well as its presence in the milk of lactating females, present potential repercussions to the developing fetus and young, naïve pups.^5,26,37,42,44,55^ Studies in non-human primates and rodents (mice and rats) exposed both in utero and in the early post-natal phase show that DA exposure causes abnormalities like repetitive behaviors and altered spontaneous behaviors; a lower seizure threshold; and structural changes to areas of the brain including the hippocampus, cortex, and amygdala.^10,21,22,38,45,55^ As such, rehabilitated neonates and pups born to dams intoxicated by DA may have a poor prognosis for survival if released or placed in permanent care. Recently a delayed manifestation of neurological disease and acute death later in life following DA exposure during development were reported in CSLs.^46^ In addition to CSL, DA can affect other species in California waters and has even been found in species as far north as Alaska, posing a potentially significant risk in the recovery of threatened species like the southern sea otter (*Enhydra lutris*) and Guadalupe fur seal (*Arctocephalus townsendi*) among others.^27,30,31,33,43^ Further studies are warranted to determine the long-term effects of DA exposure from both in utero and lactational exposure in CSL and other pinnipeds as DA exposure and strandings secondary to DA toxicosis become more frequent and expansive throughout the Pacific, and other oceans.

The NS is a useful prognostic tool as it does not require specialized equipment, is a relatively quick assessment requiring potentially only one person and is a cost-effective method that can objectively be performed without extensive training. At TMMC, a final NS <10 has been used since 2006 as a cut-off to differentiate between a releasable and non-releasable animal and the consistent use of the NS has decreased the re-strand rate for DA-intoxicated CSL over time.

When discussing the re-strand rates in the period prior to and after consistent implementation of the NS, it is also important to discuss the potential differences in the management of CSL intoxicated with DA. Although the decision-making process for management of each individual CSL is not discussed here, the differences in the proportion of released vs. euthanized CSL prior to and after implementation of the NS likely reflect changes in the process. Prior to the implementation of the NS (1998 – 2006), the proportion of CSL with DA intoxication that were released was 62.6% whereas after the implementation of the NS (2007 – 2019), the proportion of CSL with DA intoxication that were released decreased to 37.1%. Similarly, the proportion of CSL that were euthanized prior to implementation of the NS was 16% which increased to 41.3% after the NS was implemented. These differences are likely a combination of improved understanding of the disease process, differing case management styles, suspected repeat DA-exposure to CSL over time thus increasing the proportion of chronically affected CSL with generally poorer prognosis for survival after release, and the utilization of better tools to help with release assessments.

We also assessed whether the NS could be improved based on the results from the subset of 92 CSL by adjusting the relative weights of the NS sections and re-analyzing how accurate the NS was in predicting whether an animal was fit for release. By removing the “Posture” section of the NS entirely and multiplying the “Locomotion” results by three, the NS accuracy improved a small amount from 85% to 87% although the change there was no statistical support for this change.

There were limitations in assessing the NS accuracy to predict whether DA-intoxicated animals were fit for release, namely that the outcome variable (whether an animal was fit for release or not), was often determined by the same clinician treating the animal and performing the NS and the decision to release an animal was based partly on the same diagnostics used to create the NS. This introduces bias into the current study, but this was unavoidable under the circumstances and does not mean the NS is not useful as it’s a quantifiable measure of several variables that clinicians have been using to assess whether an animal is fit for release. Additionally, by analyzing variables outside of the NS such as state of nutrition at admission, whether an animal started eating while in rehabilitation, and changes in the NS final score, we aimed to help make the determination more objective for clinicians managing CSL with DA intoxication. Future clinical research could focus on determining the association of the NS with specific lesions in CSL, such as hippocampal atrophy.

While we are continually improving our understanding of DA toxicosis in marine mammals, many challenges remain, including describing and understanding long-term effects of in utero and neonatal exposure on the developing fetus and neonate, pathophysiology of the effects on organ systems outside of the central nervous and cardiac systems, and the long-term effects of repeated non-lethal exposures, among others. This study illustrates the utility of detailed clinical record keeping, and future work should focus on further refining clinical and diagnostic evaluations of naturally exposed marine mammals to clarify the pathogenesis and prognosis of DA toxicosis in these mammals.

## Acknowledgements

The authors thank the vast number of people who have cared for stranded marine mammals, contributed to the understanding of the pathophysiology of domoic acid intoxication over the past 20 years, and have improved the lives of affected marine mammals. The authors particularly thank the incredible volunteers and staff at The Marine Mammal Center and other rehabilitation facilities for their tireless efforts caring for these patients, as well our many collaborators and NOAA. Lastly, the authors thank the Dr. Jeanette Fuller Ridgway Scientific Writing Fund for providing support in the writing of this manuscript. TMMC operates under NOAA permit 18786-04.

